# Simultaneously measuring image features and resolution in live-cell STED images

**DOI:** 10.1101/190652

**Authors:** Andrew E. S. Barentine, Lena K. Schroeder, Michael Graff, David Baddeley, Joerg Bewersdorf

## Abstract

Reliable interpretation and quantification of cellular features in fluorescence microscopy requires an accurate estimate of microscope resolution. This is typically obtained by measuring the image of a non-biological proxy for a point-like object, such as a fluorescent bead. While appropriate for confocal microscopy, bead-based measurements are problematic for Stimulated Emission Depletion (STED) and similar techniques where the resolution depends critically on the choice of fluorophore and acquisition parameters. We demonstrate that for a known geometry, e.g. tubules, the resolution can be accurately measured by fitting a model that accounts for both the Point Spread Function (PSF) and the fluorophore distribution. To address the problem of coupling between tubule diameter and PSF width, we developed a technique, Nested-loop Ensemble PSF (NEP) fitting. NEP fitting enables extraction of the size of cellular features and the PSF in fixed-cell and live-cell images without relying on beads or pre-calibration. We validate our technique using fixed microtubules and apply it to measure the diameter of endoplasmic reticulum tubules in live COS-7 cells. NEP fitting has been implemented as a plugin for the PYthon Microscopy Environment (PYME), a freely available and open source software.

## Introduction

All fluorescence microscopy images distort the underlying object, failing to capture details smaller than a certain size. These distortions are described by the system’s Point-Spread Function (PSF), and knowledge of this PSF is essential when interpreting the images produced and in ensuring that quantitative measurements are accurate. For many purposes it is sufficient to summarize the effects of the PSF in a simple resolution metric, e.g. the full-width at half maximum (FWHM). A popular method for obtaining the PSF FWHM is extracting an intensity line profile from a fluorescent bead image and either directly measuring the FWHM or estimating it more accurately by fitting a Gaussian or Lorentzian (in the case of STED) model to the profile. For diffraction-limited microscopes, beads can be regarded as point-sources because they are significantly smaller than the FWHM of the PSF (beads are typically 20–100 nm compared to the ~250 nm PSF FWHM) and the fit FWHM is taken to be that of the PSF.

When considering STED microscopy, where the PSF FWHM is on the order of 50 nm, the assumption that the PSF is much larger than the bead size is no longer valid, as beads whose size is similar to that of the PSF are often needed to achieve reasonable signal levels. The resolution in STED microscopy is also strongly affected by the excitation and depletion cross-sections of the dye chosen, the choice of laser powers, and the effect of both dye micro-environment and (comparatively minor) sample-induced aberrations in the depletion beam, making beads a particularly poor proxy for the true resolution achieved when imaging cellular samples.

Measuring STED resolution on a target within the same cellular environment, labeled with the same dye(s) and imaged with the same choice of laser powers avoids most of these issues. Microtubules labeled with the same fluorescent dye as the final target structure are an attractive candidate which can be readily prepared. The simplest and most common labeling protocol that usually results in bright stainings is indirect immunofluorescence. Labeling the 25 nm outer diameter of a microtubule with primary and secondary antibodies results in a structure that is 60 nm in diameter as observed using electron microscopy (1). This is within the size range of the PSF and therefore, as with beads, the thickness of the structure is non-negligible when quantifying the resolution in a STED microscope.

To determine the impact finite object size has on resolution quantification using the popular Gaussian-and Lorentzian-fitting techniques, we simulated intensity line profiles perpendicular to the long axis of antibody-labeled microtubules imaged at various resolutions, and fit them with Gaussian and Lorentzian functions. We modeled the PSF as a Lorentzian, a common STED PSF approximation (2, 3), and the fluorophore distribution for the primary and secondary antibody labeled microtubule as an annulus of 25 nm inner diameter and 60 nm outer diameter, as measured for densely-labeled microtubules (1) (top of 1A). For most STED microscopes, the axial (z) PSF FWHM is considerably larger (500–700 nm) than the FWHM along the lateral (xy) directions. This means that the entire cross-section of a microtubule is effectively summed along the axial dimension during imaging, as shown by the red curve in figure 1A. The imaging process is simulated by convolving the fluorophore distribution with the PSF model (red curve and black curve, respectively, in fig. 1A). The resulting line profile cross section of a microtubule imaged with a 50 nm FWHM PSF is shown in figure 1A (green curve).

**Figure 1:**
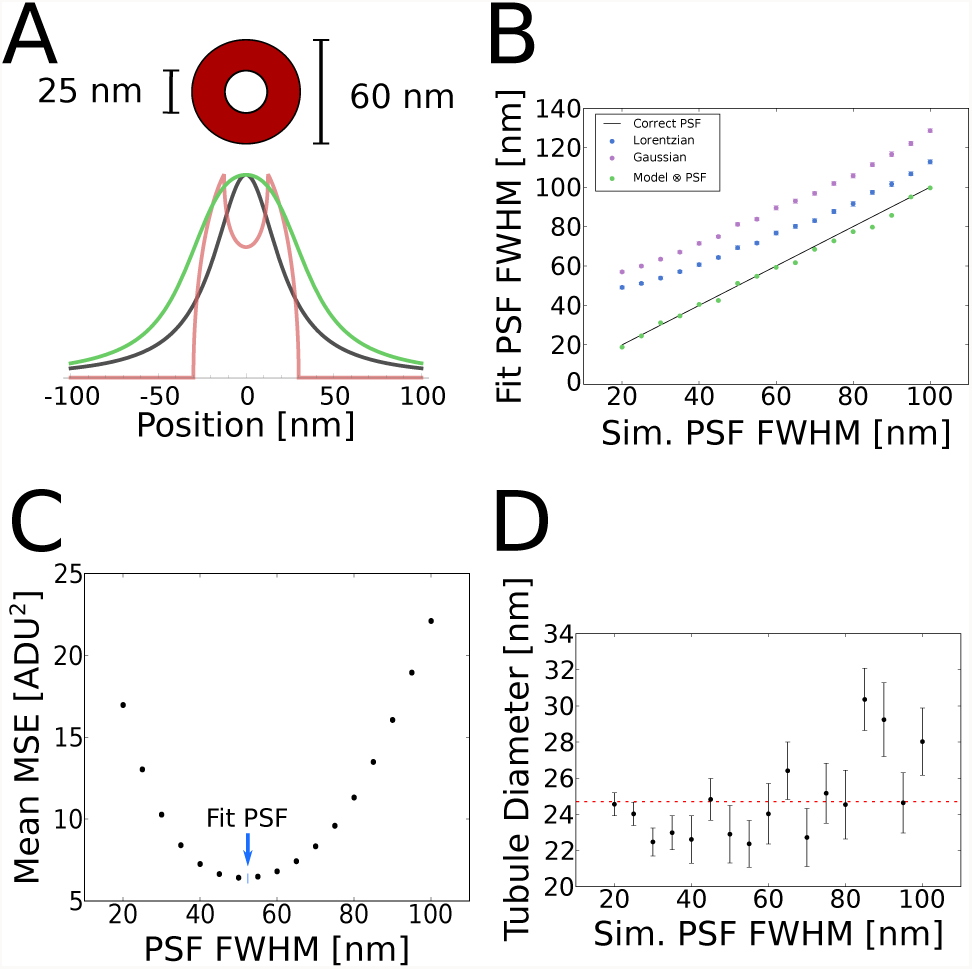
(**A**) Annulus used to model fluorophore location, where antibodies and fluorophores are bound to the 25 nm outer diameter of a microtubule (1).The red curve is a projection of the fluorophore distribution (summing over the axial dimension), the black curve is a Lorentzian function which models the PSF, and the green curve is the convolution of the red and black curves. (**B**) Microtubules were modeled and line-profiles were simulated at various resolutions, with shot noise added before being fit with simple Gaussian and Lorentzian functions. The same profiles were also fit using NEP fitting, which results in good agreement with the ground truth of the simulations. N=50 profiles were fit for each simulated PSF width. (**C**) Plot of mean MSE for fits performed with the Lorentzian-convolved model function and a specified PSF width on simulated microtubule profiles. The profiles were generated with a 50 nm PSF and added shot noise. NEP fitting performed on the same profiles minimizes the mean MSE with a PSF FWHM of 51.2 nm, as indicated by the blue arrow. (**D**) Plot of NEP fitted microtubule diameters, where images were simulated at various resolutions (N=50 profiles at each PSF width, error bars denote standard error of the mean). The ground truth diameter was 25 nm for all profiles, as shown by the dashed red line. The grey region of the plot indicates where the simulated PSF FWHM is larger than the antibodycoated tubule structure, as the tubule diameter fit is expected to be less accurate in this regime.

We simulated line profiles across microtubules imaged with STED resolutions ranging from 20 to 100 nm (PSF FWHM) and shot noise added, and fit them with simple Lorentzian and Gaussian functions. The FWHM of the Lorentzian fits were substantially larger than the PSF FWHM they were simulated with, and this effect was even more pronounced for the fitted Gaussian FWHM (fig. 1B), confirming that simple fitting of these models does not result in an accurate resolution figure. Similar Gaussian or Lorentzian fits in which the width is interpreted as the size of an imaged structure rather than resolution are popular in various types of fluorescence (superresolution) microscopy. Interpreting Figure 1B this way, shows that this approach only yields reasonable results when the PSF is much smaller (*>*3x) than the imaged structure. When used to quantify resolution, the inaccuracy increases, as expected, at higher resolutions (smaller PSF FWHM), making it particularly problematic for STED microscopy, where relative errors of 100% can easily occur. We conclude that using the FWHM of Gaussian or Lorentzian fits is an inaccurate measure for both resolution and feature size quantification.

In order to accurately determine microscope resolution or feature size from line profile cross-sections, both the microscope PSF and the geometric distribution of fluorophores on the labeled structure must be modeled in the function used for fitting. Previous efforts have fixed one of these parameters - assuming that either the PSF or the structure size is already known and enforcing this assumption either during fitting (4) or in a simulation to qualify the biased FWHM from the fit (5). These approaches are, however, limited because in biological STED microscopy both structure size and resolution are typically unknown. Fitting both of these parameters simultaneously, on the other hand, is difficult as they are not strictly independent. Increases in either parameter give rise to an increased profile width, albeit with subtly different effects on profile shape. At the signal-to-noise level typical of a single profile it is difficult to separate the effects of the two parameters. This coupling can result in inaccurate estimates for both values. Here, we present a tool which overcomes this challenge, and allows simultaneous fitting of structure size and PSF width.

## Ensemble PSF Fitting

Treating both the PSF width and the structure size as free parameters, it should, in principle, be possible to determine both simultaneously. To make the fit more robust, we fit multiple profiles and exploit the prior knowledge that the PSF width should be the same for each profile. We accomplish this by performing a two-layer nested fit, such that in the inner fit, all tubules are fit with the PSF FWHM constrained to be the same value, *γ*, and the mean squared error (MSE) for each tubule is reported. The outer fit is then responsible for finding the value of *γ* which minimizes the mean MSE taken over all of the tubule fits, which has a propensity to be well-behaved and smooth over values near the expected PSF FWHM (fig. 1C). This technique, which we refer to as nested-loop ensemble PSF (NEP) fitting, constrains the fit enough that more accurate PSF widths and microtubule diameters can be determined, as shown in figure 1B, D.

NEP fitting using the antibody-coated tubule model yielded significantly better results than fitting with plain Gaussian or Lorentzian functions, and the PSF widths calculated by the fit are in close agreement with the ground truth (fig. 1B). Notably, NEP fitting with the antibody-coated tubule model simultaneously yields accurate measures of the simulated microtubule diameter, 25 nm, for all simulated PSFs with FWHM equal to or less than 60 nm, which is the value at which the PSF FWHM becomes larger than the outer diameter of the antibody coat (fig. 1D). For structures whose size does not vary in a cell, e.g. microtubule diameters, the structure size can additionally be constrained as an ensemble parameter during the fit, although we did not find this to be a necessary step.

## Software and Validation

We implemented NEP fitting for STED images of label-filled or surface-labeled tubules in the PYthon Microscopy Environment (PYME). Line profiles of a user-defined thickness are extracted from images loaded into PYME, after which they are fit using a variety of model functions. Alternatively, they can be saved/appended to two file formats (HDF, json) for later analysis or ensemble fitting with profiles from multiple images. The line profile extraction GUI is shown in figure 2A.

**Figure 2:.**
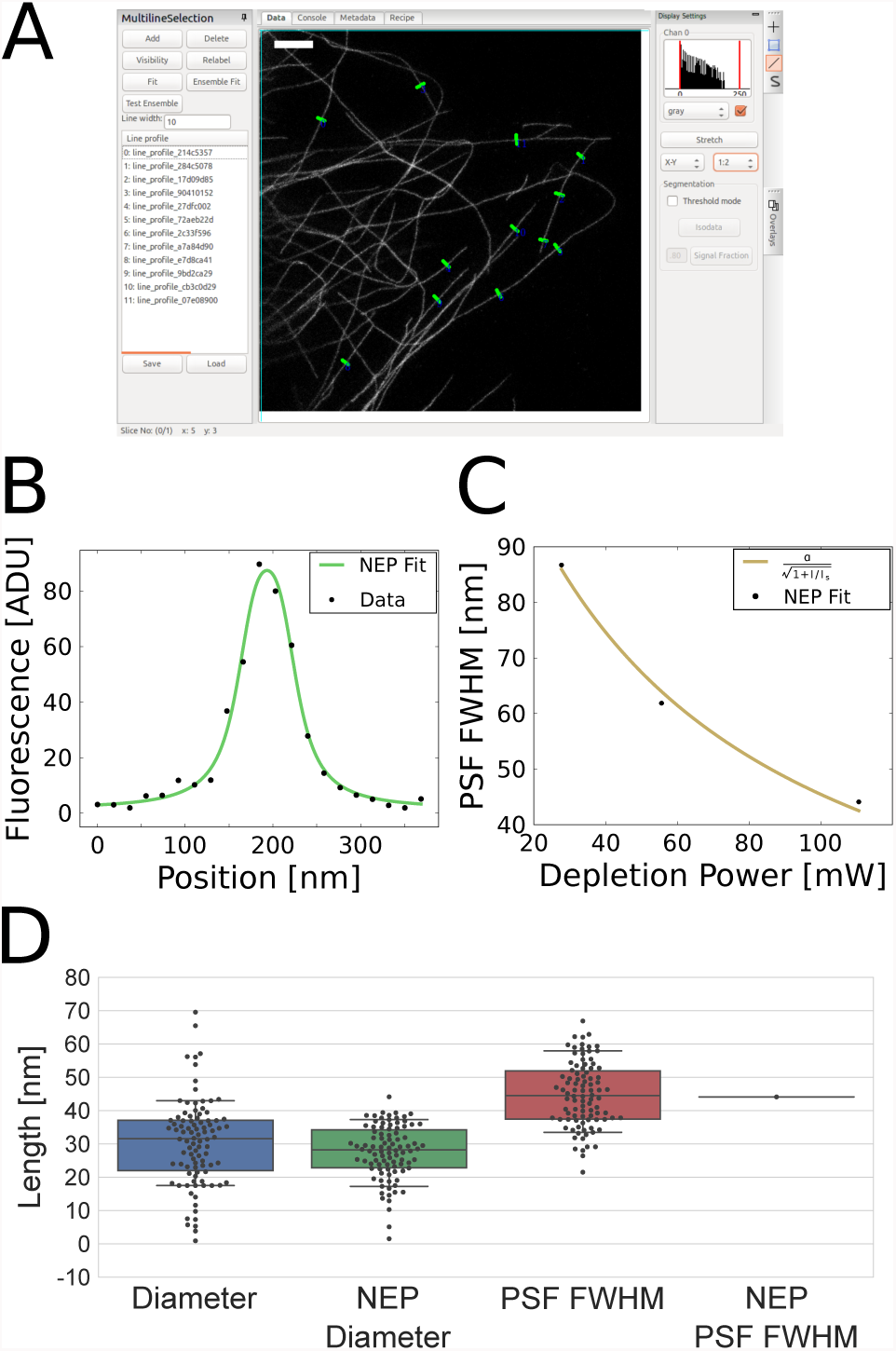
(**A**) PYME GUI showing a STED image of immunolabeled microtubules in a COS-7 cell, imaged with 110.6 mW STED laser power. Green lines show user-selected profiles to be used for fitting. (**B**) Plot of raw data and NEP fit of a microtubule profile from (*A*).(**C**) Plot of NEP-fitted PSF widths from STED images of microtubules acquired with different STED powers, which scales as expected by theory (n=74, n=71, and n=94 profiles extracted from N=8, N=8, and N=12 images of N=3, N=3, and N=6 cells, acquired at 27.7, 55.6, and 110.6 mW STED laser powers, respectively). (**D**) Swarmand boxplots of microtubule diameters and PSF FWHM values determined using NEP fitting, where the PSF is constrained to be the same for all microtubule line profile cross sections, and without NEP fitting (standard least-squares fitting), where the PSF is varied separately for each tubule fit.

STED PSF size is dependent on the STED laser power with a scaling of 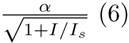 (6). To test the efficacy of ensemble fitting on real data, we imaged primary and secondary antibody labeled microtubules using a Leica SP8 STED 3X microscope with different STED laser powers. After extracting 239 line profiles from a total of 28 images (example shown in fig. 2B, n=74, n=71, and n=94 profiles extracted from N=8, N=8, and N=12 images of N=3, N=3, and N=6 cells, acquired at 27.7, 55.6, and 110.6 mW STED laser powers, respectively), NEP fitting was performed. As shown in figure 2C, this demonstrates that our fit is responsive to changes in PSF size resulting from varied STED laser powers, and can reproduce the expected scaling of the PSF widths with the STED laser power.

To determine if our ensemble (NEP) fitting approach in which the PSF width is constrained to a single, global value for all profiles is beneficial, we compared its results with those obtained when both PSF width and tubule diameter were optimized on a per-profile basis. We performed this comparison on images recorded with 110.6 mW depletion power. While fits performed without an ensemble PSF have more degrees of freedom and therefore usually yield smaller residuals, they often come at the expense of accuracy in the measured values. This can be seen in figure 2D, where the standard least squares fit results in an average microtubule diameter of 30 ± 13 nm (mean ± SD), compared to the NEP-fitted value of 28 ± 8 nm (mean ± SD), and the expected value of 25 nm. NEP fitting, with its global PSF constraint, indeed improves the measurement of the microtubule diameter, as evidenced by the reduced spread in tubule diameters and average value closer to the expected 25 nm. The PSF FWHM of standard least squares fitting was 45 ± 10 nm (mean ± SD), compared to the NEP-fitted value of 44 nm. We note that fitting a single line profile would not provide a reliable measure of either PSF FWHM or tubule diameter, and could lead to relative errors of 100% for both values. The minimum number of profiles necessary for robust NEP fitting depends on the fluorophore distribution and the relative PSF size. However, even for cases of low signal-to-noise ratio (SNR), we found 100 profiles to be sufficient for the fluorophore distributions tested (see fig. S1, S2).

## Application to live-cell images

While the robust PSF measurement in fixed cells by NEP fitting is a substantial improvement over bead calibrations, a large advantage of NEP fitting is that it can be performed on live-cell data for *in-situ* resolution calibration in the most biologically relevant state. We applied ensemble PSF fitting in live-cell STED images of label-filled or surface-labeled endoplasmic reticulum (ER) tubules, using ss-SNAPKDEL or SNAP-Sec61*β*, respectively (fig. 3A-D). In order to fit the label-filled tubules, we modeled the fluorophore distribution perpendicular to the long axis of the tubule as a filled circle, which projects as 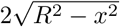 where *R* is the radius, which we then convolved with a Lorentzian to account for the imaging process. We modeled the fluorophore distribution for surface-labeled tubules as an annulus, like the antibody-labeled microtubules only with a thinner coat. SNAP-tag (7) is about 4 nm in diameter and the organic dye itself can be estimated to have a radius of 0.5 nm assuming they are both globular (8), resulting in a 4.5 nm thick annulus. Fits of ER tubule diameter for various test PSF widths show stronger coupling between the tubule diameter and PSF width for label-filled tubules than surface-labeled tubules (fig. 3E,F). The PSF widths from the NEP fits were similar for label-filled and surface-labeled profiles (45.8 nm and 43.7 nm FWHM, respectively) as shown in figure 3G, H.

**Figure 3:**
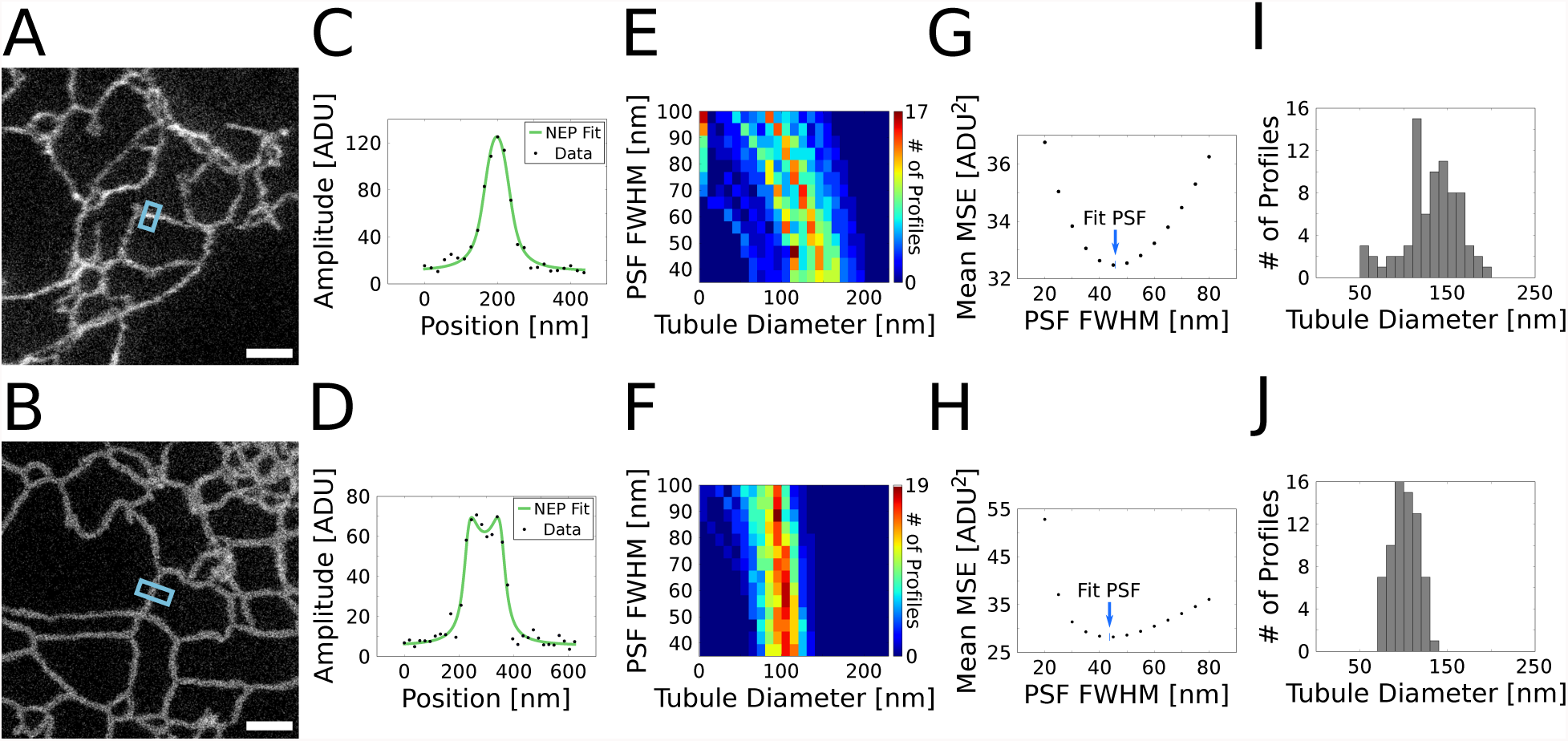
(**A**, **B**) Live-cell STED images of label-filled ((*A*), ss-SNAP-KDEL) and surface-labeled ((*B*), SNAP-sec61*β*) ER. (**C**, **D**) Fluorescence line profiles, averaged over 10 pixels along the long axis of the tubule, extracted from (*A*) and (*B*) (black points), respectively, and fit using NEP fitting, where the PSF width is constrained to be the same for all line profiles. (**E**, **F**) Heatmaps showing the coupling between tubule diameter and PSF FWHM when standard least-squares fitting is performed with varied (fixed) PSF FWHM. Line profiles of ER with label-filled (*E*, ss-SNAP-KDEL) and surface-labeled (*F*, SNAP-sec61*β*) tubules were fit with the PSF FWHM fixed to a value that was iteratively changed. Intensity corresponds to number of profiles (n=77 and n=69 profiles were extracted from N=7 and N=7 STED images of N=4 and N=2 cells, acquired for ss-SNAP-KDEL and SNAP-sec61*β* expressing cells, respectively. Images corresponding to each label were acquired on a single day.) (**G**, **H**) Mean MSE where the mean is taken over all label-filled (*G*) and surface-labeled (*H*) ER tubule fits which were performed with the PSF FWHM fixed to a value which was iteratively changed. The blue arrow indicates the PSF FWHM found by performing NEP fitting on the same tubule line profiles. (**I**, **J**) Label-filled (*I*) and surface-labeled (*J*) ER tubule diameters fit with the PSF estimated using ensemble PSF fitting (45.8 nm PSF FWHM for ss-SNAP-KDEL, 43.7 nm PSF FWHM for SNAP-sec61*β*). The mean and standard deviations were 132 ± 30 nm and 101 ± 15 nm for ss-SNAP-KDEL and SNAP-sec61*β*, respectively.

Notably, the standard deviation for both the label-filled and surface-labeled tubule diameters is fairly large: 30 and 15 nm, respectively (with mean values of 132 nm and 101 nm). To test whether this variability in tubule diameter is primarily biological in nature, or dominated by the SNR-limited fit precision, we simulated tubule profiles of known diameter with similar signal-to-noise ratios, convolved with 50 nm FWHM Lorentzians. The distributions of fitted diameters were more narrow than those observed in the live-cell images, with standard deviations of only 5 nm for the membrane-labeled tubules, and 12 nm for the label-filled tubules (Fig. S1). The larger range of label-filled tubule fitted diameters is expected because the fluorophore distribution orthogonal to the long axis of the tubule looks more similar to the PSF than a surface-labeled fluorophore distribution. This is reflected in the live-cell data, where the tubule diameter is more strongly coupled to the PSF width for the label-filled tubules, as shown in the heat-maps of tubule diameter histograms when fit with various fixed PSF widths (Figures 3E,F). We therefore expect roughly half of the spread in tubule diameter to be biological in origin. Notably, the NEP fitting estimates for tubule diameter and PSF width do not suffer from systematic errors when the signal-to-noise ratio (SNR) of the profile is decreased. However, the variability in fit tubule diameters for individual profiles is increased for lower SNR profiles (Fig. S2).

## Discussion

Traditional methods of resolution calibration in STED microscopy are problematic for biological quantification. The NEP fitting method introduced in this paper addresses this issue and provides a robust and practical means to both quantify the performance of a microscope, as well as improve feature measurements within the image. Its implementation in a freely-downloadable, open source, cross-platform software package allows for rapid adoption by others, without requiring mathematical or programming expertise. The principle of ensemble fitting can be readily extended to other fluorescence microscopy modalities, e.g. confocal, by substituting a different functional representation of the PSF when producing the model function for fitting; the only requirement is that the labeling geometry of the structure is known. This known geometry is not limited to tubules, and can be extended to fit objects like beads or vesicles, which would be useful for cell-trafficking studies. The accurate measure of PSF width afforded by NEP fitting can be used to quantify microscope performance under various conditions, refine models of organelle morphology, and remove uncertainty in parameter selection for deconvolution or other image enhancement algorithms.

## Software Availability

All line profiles were drawn, extracted, and fit using the open source PYthon Microscopy Environment (PYME) and the NEP Fitting plug-in, which are both freely available (9, 10).

## Acknowledgements

This work is supported in part by NIH grant S10 OD020142 for imaging resources (for the Leica TCS SP8 STED 3X microscope), the G. Harold & Leila Y. Mathers Foundation, the Wellcome Trust (095927/A/11/Z, 203285/B/16/Z), and the Yale Diabetes Research Center (NIH P30 DK045735). A. E. S. B. acknowledges support by an NIH training grant (T32 GM008283). J. B. discloses significant financial interest in Bruker Corp. and Hamamatsu Photonics. We thank Phylicia Kidd and Mark Lessard for providing biological test samples, and Zach Marin for helpful discussions.

## Supporting References

References (11–14) appear in the Supporting Material.

